# scHiCSRS: A Self-Representation Smoothing Method with Gaussian Mixture Model for Imputing single cell Hi-C Data

**DOI:** 10.1101/2021.11.09.467824

**Authors:** Qing Xie, Shili Lin

## Abstract

**Motivation:** Single cell Hi-C techniques make it possible to study cell-to-cell variability in genomic features. However, excess zeros are commonly seen in single cell Hi-C (scHi-C) data, making scHi-C matrices extremely sparse and bringing extra difficulties in downstream analysis. The observed zeros are a combination of two events: structural zeros for which the loci never interact due to underlying biological mechanisms, and dropouts or sampling zeros where the two loci interact but are not captured due to insufficient sequencing depth. Although quality improvement approaches have been proposed as an intermediate step for analyzing scHi-C data, little has been done to address these two types of zeros. We believe that differentiating between structural zeros and dropouts would benefit downstream analysis such as clustering.

**Results:** We propose scHiCSRS, a self-representation smoothing method that improves the data quality, and a Gaussian mixture model that identifies structural zeros among observed zeros. scHiCSRS not only takes spatial dependencies of a scHi-C 2D data structure into account but also borrows information from similar single cells. Through an extensive set of simulation studies, we demonstrate the ability of scHiCSRS for identifying structural zeros with high sensitivity and for accurate imputation of dropout values in sampling zeros. Downstream analysis for three real datasets show that data improved from scHiCSRS yield more accurate clustering of cells than simply using observed data or improved data from several comparison methods.

**Availability and Implementation:** The scHiCSRS R package, together with the processed real and simulated data used in this study, are available on Github at https://github.com/sl-lin/scHiCSRS.git.

**Supplementary information:** Supplementary data are available online.

## 1 Introduction

The spatial organization of chromosomes in a cell nucleus is not random; rather, it is dynamic and closely linked to genome functions and disease mechanisms (Dekker, 2008). Harnessing the power of next-generation sequencing technologies, the Hi-C technology enables a high resolution, genome-wide three-dimensional (3D) view of the chromosomal organization (Lieberman-Aiden et al., 2009), and it has been applied to analyze different types of cells (Rao et al., 2014; Kim et al., 2017; Darrow et al., 2016). The original Hi-C technique produces bulk data, averaging chromosome conformation over millions of cells and resulting in limited information on cell-to-cell variability (Fraser et al., 2015). Recent single cell Hi-C assays, on the other hand, enable the analysis of whole-genome structures for single cells (Nagano et al., 2013) and has the potential to identify rare cell populations or cell sub-types in a heterogeneous population (Ramani et al., 2019)

Interpreting single cell Hi-C (scHi-C) data is challenging because of data sparsity (observed zeros) and low sequencing depth (Nagano et al., 2015). Due to the increase of data dimension, the coverage of scHi-C (0.25 − 1%) is much smaller than that of RNA-seq (5 − 10%) (Zhou et al., 2019), leading to additional difficulty for analyzing scHi-C data. The observed zeros are a mixture of two types of events: some are structural zeros because the pairs do not interact with each other due to the underlying biological mechanisms, while others are dropouts or called sampling zeros as a result of low sequencing depth. While dropouts happen at random, structural zeros do not. Differentiating between structural zeros and dropouts and imputing the latter can lead to improved downstream analyses such as clustering and 3D structure inference.

The zero-inflated phenomenon is also observed in single cell RNA (scRNA) research. Currently, there is considerable research on imputation for scRNA data, with the concept of structural zero well defined and inferences made to distinguish structural zeros and dropouts (van Dijk et al., 2017; Chen et al., 2018; Li and Li, 2018; Mongia et al., 2019; Peng et al., 2019; Hu et al., 2020; Zhou et al., 2020; Zand and Ruan, 2020; Rao et al., 2021). In contrast, the concept and inference of structural zeros and dropouts have not been widely discussed in scHi-C research, although we note that in several papers that aim to assess data reproducibility (Yang et al., 2017; Ursu et al., 2018), construct 3D structures (Zhu and Wang, 2019), or cluster single cells (Zhou et al., 2019), imputing values for observed zeros has been treated as an intermediate data enhancing step. In a recent contribution, we explored the potential of using scRNA methods for analyzing scHi-C data and achieved some success (Han et al., 2020). However, the issue of scRNA methods not accounting for spatial correlation – a hallmark of Hi-C data – was also identified.

In the Hi-C literature for quality improvement for bulk or single cells data, kernel smooth, random walk, and convolutional neural network are the main ideas (Yang et al., 2017; Ursu et al., 2018; Zhou et al., 2019; Zhu and Wang, 2019). The 2D mean filter approach (a kernel smoothing method) directly replaces each cell of a 2D contact matrix with the mean count of all contacts in its genomic neighborhood. For example, HiCRep (Yang et al., 2017) applies such a filter to assess the reproducibility of Hi-C data. ScHiC-Rep (Zhen et al., 2021) applies a uniform kernel to cluster scHi-C data. scHiCluster (Zhou et al., 2019) applies a convolution-based imputation including a mean filter to help cluster cells. Different from a 2D mean filter that takes an average of the genomic neighbors, kernel smooth uses a weighted average of neighboring observed counts. The weight is defined by a kernel, which gives more weight to closer genomic neighbors. For instance, SCL (Zhu and Wang, 2019) applies a 2D Gaussian function to impute scHi-C contact matrices and further infers the 3D chromosome structures from the enhanced Hi-C data. GenomeDISCO (Ursu et al., 2018), on the other hand, uses a random walk on the contact map to “smooth” the observed counts, and it shows that taking three steps of the random walk would lead to the best results in general. scHiCluster (Zhou et al., 2019) also uses the idea of a random walk, but with restarts, to capture the topological structure. Convolutional neural network is also an approach commonly applied to infer a high-resolution Hi-C matrix from a low-resolution one. HiCPlus (Zhang et al., 2018) and DeepHiC (Hong et al., 2020) are examples of such supervised learning techniques.

Although taking spatial correlation in a 2D data matrix into consideration, the current methods as discussed above enhance each Hi-C data matrix independently without considering other information, such as data from similar cells. Further, inference on structural zeros and dropouts is rarely discussed, although the identification of such may play an important role in downstream analyses. In an attempt to make fuller usage of available information and to distinguish structural zeros from dropouts, in this paper, we develop scHiCSRS, a self-representation smoothing method. It not only borrows information from 2D neighborhoods but also takes similar single cells into account. Further, as part of the scHiCSRS package, we propose a Gaussian mixture model to separate the zeros into structural zeros and dropouts. Through an extensive set of simulation studies and real data analyses, we showed that scHiCSRS can accurately identify structural zeros and impute the dropouts. We also compared scHiCSRS with other methods for data quality improvement and downstream clustering analyses.

## 2 Materials and Methods

The overall goal of scHiCSRS is to enhance scHi-C data and make inference on structural zeros (Figure 1). scHiCSRS takes spatial dependencies of scHi-C 2D data structure into consideration while also borrows information from similar single cells. scHiCSRS was motivated by scTSSR (Jin et al., 2020) that recovers scRNA data using a two-sided sparse self-representation method, but there are two major differences. Firstly, scTSSR uses the expression of all genes in the same cell while scHiCSRS only considers counts in a 2D matrix neighborhood, which helps capture local dependencies (Zhen et al., 2021). Secondly, scTSSR has an interaction term that involves elements in the same row and column; however, scHiCSRS does not include such a term because other positions in other single cells should have no direct influence on the position to be imputed. Based on the quality-improved data, we further apply a Gaussian mixture model to identify structural zeros.

**Figure 1:**
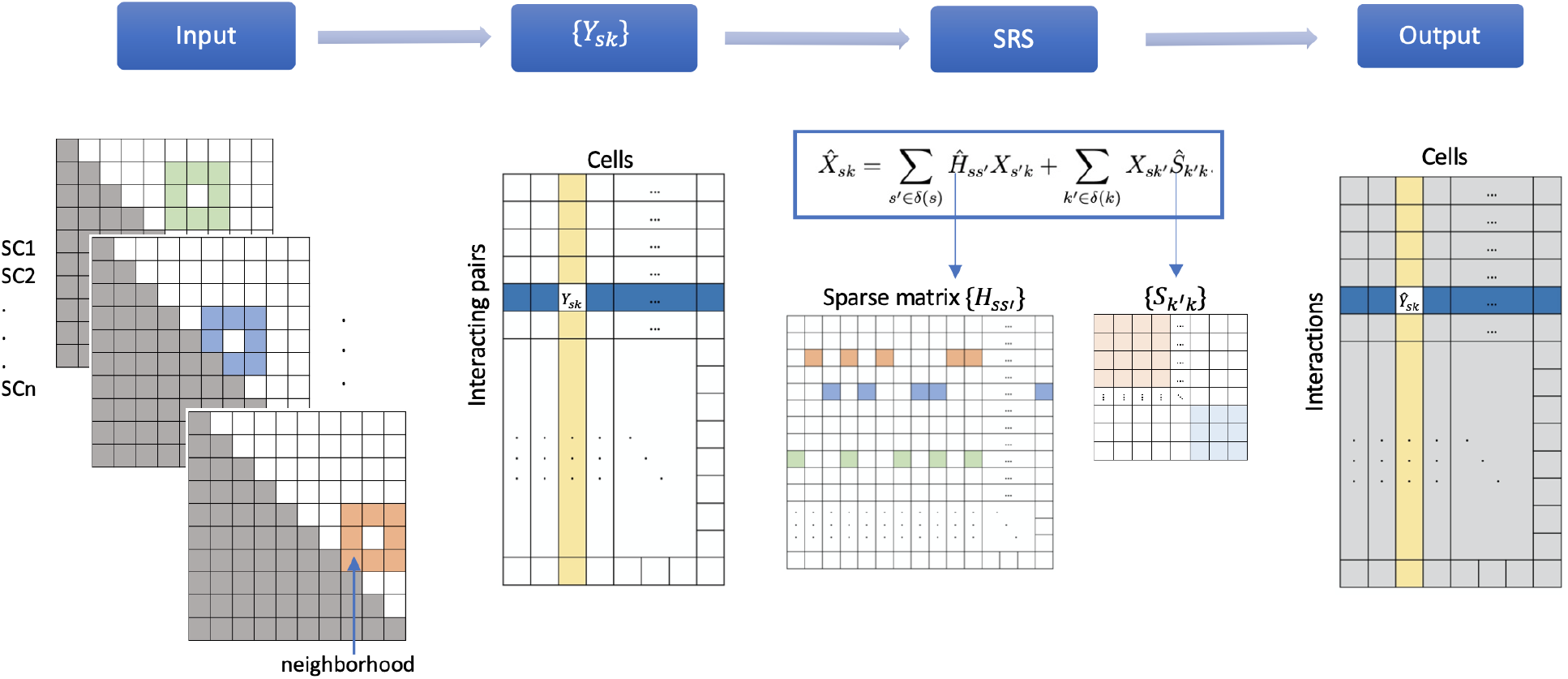
Schematic of the scHiCSRS algorithm. *Input:* the input includes multiple scHi-C contact matrices, with the colored region of a cell denoting the neighborhood of a position enclosed. *Data matrix* {*Y*_*sk*_} : the single cells are organized into a big matrix, with each row representing a pair of interacting loci and each column being the upper triangular of a single cell contact matrix. *SRS:* A self-representation model is used to enhance the entries in the matrix *X* (normalized from the observed matrix *Y*); since SRS only borrows information from 2D neighborhoods, the coefficient matrix *H* is sparse with its values in most positions (not in the neighborhood of a position) set to 0; if the input single cells are composed of more than one type, the matrix *S* is also sparse, with only non-zero blocks along the diagonal because we only consider the influence from similar single cells. *Output:* the output is the enhanced matrix 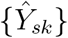, based on which we can perform additional analyses.

### 2.1 Self-representation smoothing model

Suppose we have contact matrices for *K* single cells. Let *Y*_*ijk*_ represents the observed interaction frequency between loci *i* and *j* (*i* ≤ *j*) for single cell *k* (*k* = 1, … , *K*), where a locus is a gnomic segment and {*Y*_*ijk*_}_*n*×*n*_ is a symmetric 2D matrix of dimension *n* × *n* for each single cell *k*, 1 ≤ *k* ≤ *K*, where *K* is the number of single cells and *n* is the number of genomic loci considered. We combine the 2D contact matrices of all single cells into a big matrix {*Y*_*sk*_} (*s* = 1, … , *N* = *n*(*n* + 1)/2, *k* = 1, … , *K*) of dimension *N* × *K* with each column being the upper triangular of a single cell 2D matrix. We first normalize each cell so that all cells have the same sequencing depth (the median—med—across all cells), then we log-transform the normalized matrix as follows:

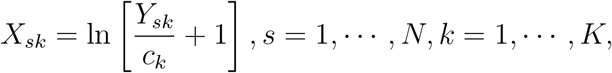

where *c*_*k*_ = Σ_*s*_ *Y*_*sk*_/med{Σ_*s*_ *Y*_*sk*_, *k* = 1, … , *K*} is the depth-adjusted normalization factor for cell *k*, and a pseudo count of 1 is added due to the existance of observed zeros.

For each *X*_*sk*_, there are two types of information that we use for the smoothing process: the neighborhood *δ*(*s*) and the collection of similar cells *δ*(*k*) at the same position; that is, *δ*(*s*) contains the 2D neighbors of position *s* (but not *s* itself) while *δ*(*k*) contains all the cells that are similar to *k* (but not *k* itself). To smooth the contact matrix, we assume that the contact count of each pair is a linear combination of these two types of information. Therefore, we propose the following self-representation smoothing (SRS) model for obtaining a “smoothed” scHi-C matrix:

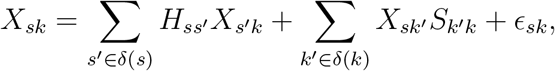

where the {*H*_*ss’*_}_*N*×*N*_ and the {*S*_*k’ k*_}_*K*×*K*_ matrices are described in the following.

For convenience, the neighborhood *δ*(*s*) is taken to be a regular one, as shown in Figure 1, although the size and shape may be modified as appropriate. For all the data analyses carried out in this paper, we use a regular neighborhood with 24 neighbors. The *N* × *N* matrix {*H*_*ss’*_}_*N*×*N*_ describes the influence of neighbor *s*’ on position *s* so that only positions within the neighborhood have a positive coefficient and the others are set to 0, leading to a sparse matrix (Figure 1). The *K* × *K* matrix {*S*_*k’k*_}_*K*×*K*_ describes the influence of cell *k*’ on cell *k* and is set in such a way that only similar cells *k*’ ∈ *δ*(*k*) have a positive influence, the rest is set to 0. Thus, if the input single cells are of different types, the matrix *S*_*k’ k*_ would have non-zero blocks along the diagonal with each block being the coefficients for single cells of the same type ( Figure 1). Descriptions on how to obtain the estimates of the coefficient matrices {*H*_*ss*’_} and {*S*_*k’ k*_} are provided in the supplementary material. Once we obtain their estimates, denote as 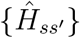 and 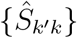, respectively, the imputed value is calculated as

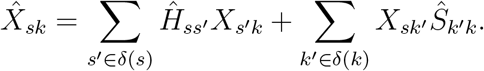

Although the imputed value 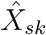 borrows information from the contacts in neighboring positions in the same cell and other cells at the same position, it does not take the observed value *X*_*sk*_ itself into consideration directly. Therefore, we couple the above procedure with the idea of a Bayesian model for scRNA data (Huang et al., 2017). We model the observed count *Y*_*sk*_ (without normalization or log-transform) as follows: *Y*_*sk*_ ~ *Poisson*(*c*_*k*_*λ*_*sk*_) and *λ*_*sk*_ ~ *Gamma*(*α*_*sk*_, *β*_*sk*_), where *λ*_*sk*_ represents the normalized (med) true interaction intensity and *c*_*k*_ is the normalization factor as defined above. The marginal distribution of *Y*_*sk*_ is then a negative binomial, allowing for over-dispersion. The prior mean (for the Gamma distribution at the normalized scale) is set to be 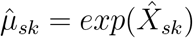 and the prior variance is estimated through a constant noise model across all cells. Reparameterization leads to the estimated shape and rate parameters, 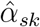 and 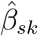. The posterior distribution is then 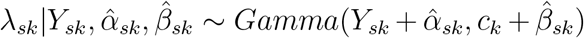. We use the posterior mean to estimate *λ*_*sk*_ as follows:

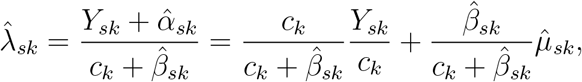

which is a weighted average of the normalized observed contact counts and the prior mean estimated from SRS. The final imputed value for *Y*_*sk*_, in the original scale, is 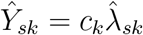

### 2.2 Gaussian mixture model

Since the self-representation smoothing model does not have an internal mechanism for separating structural zeros from dropouts, we further propose a Gaussian mixture model on the imputed data 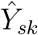 to address this issue. We start by normalizing the imputed matrix to the median library size and taking the log_10_ transformation with a pseudo count 1, the same as described in section 2.1, albeit it is now with the imputed, not the raw counts.

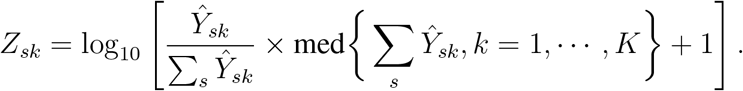

Without loss of generality, we assume all the cells are of the same type so that we can use the notation already defined above. If there are multiple known types, then the Gaussian mixture model will be applied to each separately. For a pair of loci (i.e. a position in the 2D Hi-C data matrix) that has zero interaction counts in all the single cells, they are automatically labeled as structural zeros without being subjected to the mixture analysis. For the remaining pairs with zeros in some cells and nonzeros in other cells, collectively denoted as 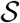, we assume that

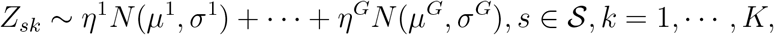

where 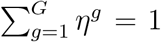 and *μ*^1^ < *μ*^2^ < … < *μ*^*G*^. That is, the imputed values at the positions with observed zero in some cells follow a *G*-component Normal mixture distribution. For a position in a cell that has high imputed interaction frequencies, captured by a component with a higher mean, an observed zero is more likely a dropout; whereas if the imputed interaction frequency is low, captured by a component with a lower mean, then an observed zero may be a true structural zero. The parameters are estimated using the Expectation-Maximization (EM) algorithm for a given *G*, and the best *G*, 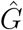, is selected based on BIC (Claeskens et al., 2008). We then calculated 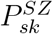, the probability of being structural zero for each position 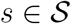 in each single cell *k* as follows:

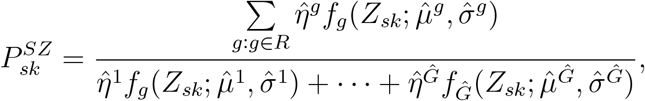

where the *f*’s are the Normal density functions, and *R* is the Gaussian components designated as the structural zero component(s) based on the following rule. If 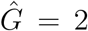, the first component is chosen to capture structural zeros. If 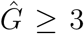, denote the distances between adjacent means to be 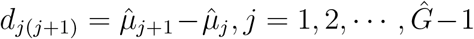. If *ξd*_12_ ≤ *d*_23_ for a large multiple *ξ* (say, *ξ* = 10), meaning that the first two components are close to each other but are far away from the third component, we choose the first and second as structural zero components; otherwise, only the first component is treated as capturing structural zeros. If both of the first and second components are already chosen as capturing structural zeros, we continue the process using the same criterion to ascertain whether additional successive components, up to 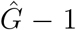, should be chosen. Finally, an observed zero is classified as a structural zero if 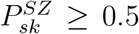, although other threshold values may also be considered.

### 2.3 Performance evaluation criteria

We evaluate the performance of scHiCSRS and compare it with other data quality improvement methods by considering several criteria. First, we evaluate the ability of scHiCSRS to identify structural zeros among the observed zeros, and to compare its performance with methods in the literature. Specifically, for the comparison methods, since they do not have an internal mechanism for identifying structural zeros, we label an observed zero as a structural zero if the imputed value is less than 0.5, following suggestions in the literature (Han et al., 2020). To measure the ability of a method (scHiCSRS or a comparison method) to separate structural zeros from sampling zeros, we call the proportion of true structural zeros identified as the *power* or *sensitivity*, defined as the proportion of underlying structural zeros correctly identified. Similarly, we call the proportion of true dropouts, defined as the proportion of underlying sampling zeros correctly identified, as the *specificity* to measure the ability of a method for correctly identifying dropouts. Since the identification of structural zeros and dropouts depends on the decision rules (a threshold on the probability for the Gaussian mixture model or a threshold on the imputed value for the comparison methods), we also explore a range of thresholds, with the result measured as the area under the curve (AUC) – the curve being the conventional receiver operating characteristic (ROC) curve – for a more thorough comparison of methods. We use the absolute errors between the imputed and the expected values to further assess the imputation accuracy of scHiCSRS and the comparison methods. Additionally, we use the correlation between the imputed and the expected to measure the aggregate performance of a method.

## 3 Simulation study

### 3.1 Data generation

To mimic real data, we use three 3D structures on a segments of chromosome 1 (the first 61 mega bases loci) recapitulated using SIMBA3D (Rosenthal et al., 2019) from three K562 single cell Hi-C 2D matrices (Flyamer et al., 2017). For each structure (single cell), based on the estimated 3D coordinates (*x*_*i*_, *y*_*i*_, *z*_*i*_) (1 ≤ *i* ≤ 61), we firstly generate the interaction intensity matrix *λ* = {*λ*_*ij*_} with the following model:

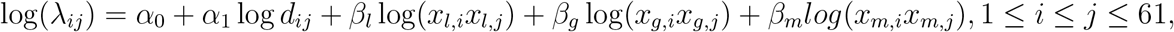

where 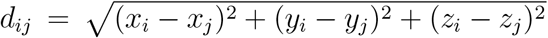 is the distance between loci *i* and *j*; *x*_*l,i*_ ~ Unif(0.2, 0.3), *x*_*g,i*_ ~ Unif(0.4, 0.5), and *x*_*m,i*_ ~ Uni*f* (0.9, 1) mimic covariates such as fragment length, GC content, and mappability score (Park and Lin, 2019), and *β*_*l*_, *β*_*g*_, and *β*_*m*_ are the corre-sponding coefficients of the covariate terms; *α*_1_ is set to −1 following the typical biophysical model; and *α*_0_ is a scale parameter that we used to control sequencing depths.

These three structures are designated as three “types” (I, II, and III) of single cells. For each type, we simulate *n* single cells, with varying numbers of *n*’s as described below. To simulate sparse 2D matrices with both structural zeros and dropouts, we define a threshold *b* as the lower 10% quantile of the *λ*_*ij*_’s. For those *λ*_*ij*_ < *b*, we randomly select half of them to be structural zeros candidates; among them, 80% are randomly selected to be structural zeros across all *n* single cells. For a particular single cell, we randomly select half of the remaining 20% candidates to be structural zeros. For those candidates that are selected as structural zeros, their new *λ*_*ij*_ are set to be zero while for those that are not selected to be structural zeros, the *λ*_*ij*_ values are left unchanged in the original *λ* matrix. This procedure makes each single cell has its specific *λ*\* matrix (containing “expected” values). Based on the *λ*\* matrix, we generate the contact counts using a Poisson distribution with the intensity parameter being the corresponding 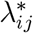 for a particular single cell. This step also produces dropouts that are observed zeros but their underlying true values are nonzero. Using three sets of parameters (Table S1), we simulated single cells for type I, II, and III with three sequencing depths (7k, 4k, and 2k) and three sample sizes of cells (10, 50, 100).

### 3.2 Results

We choose three smoothing methods that have been used as an intermediate step to enhance Hi-C data for comparison with scHiCSRS. These three methods are mean filter ( MF) as in HiCRep (Yang et al., 2017), which replaces each contact with the average count of its neighborhood region, Gaussian kernel smooth (GK) as in SCL (Zhu and Wang, 2019), which uses a weighted average of neighboring observed data and the weights determined by a Gaussian kernel, and random walk (RW) as in GenomeDISCO (Ursu et al., 2018), which takes a 3-step random walk.

For correct identification of structural zeros, scHiCSRS has a power of near 0.9 or higher in all situations (Figure 2(a) and Table S2). In contrast, the performance of the three comparison methods fluctuates greatly with sequencing depth: it may be as high as 0.85 when the sequencing depth is 2k, but may be down to zero when the sequencing depth is 7k. We also used ROC curves to explore the interplay between correct identification of structural zeros and dropouts for a fair comparison of all methods; the AUC for scHiCSRS is much higher than the comparison methods (Figure 2(b) and Table S3).

**Figure 2:**
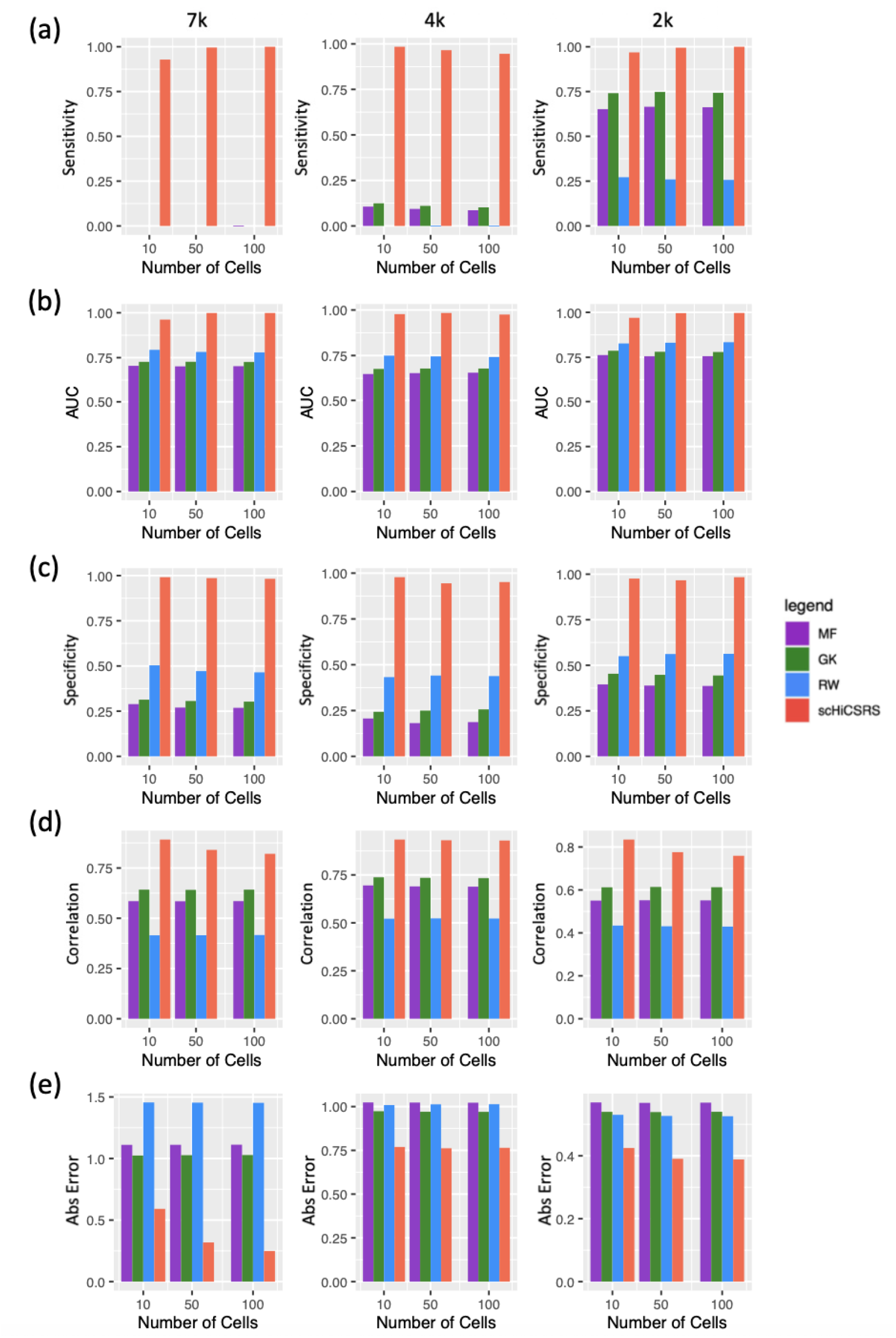
Barplots of several criterion values for type I cells over three sequencing depths: 7k (1st column), 4k (2nd column), and 2k (3rd column): (a) sensitivity for detecting structural zeros; (b) areas under ROC curves constructed with a range of thresholds; (c) specificity for identifying dropouts; (d) correlation between imputed values and expected; and (e) absolute difference between imputed and expected.

**Figure 3:**
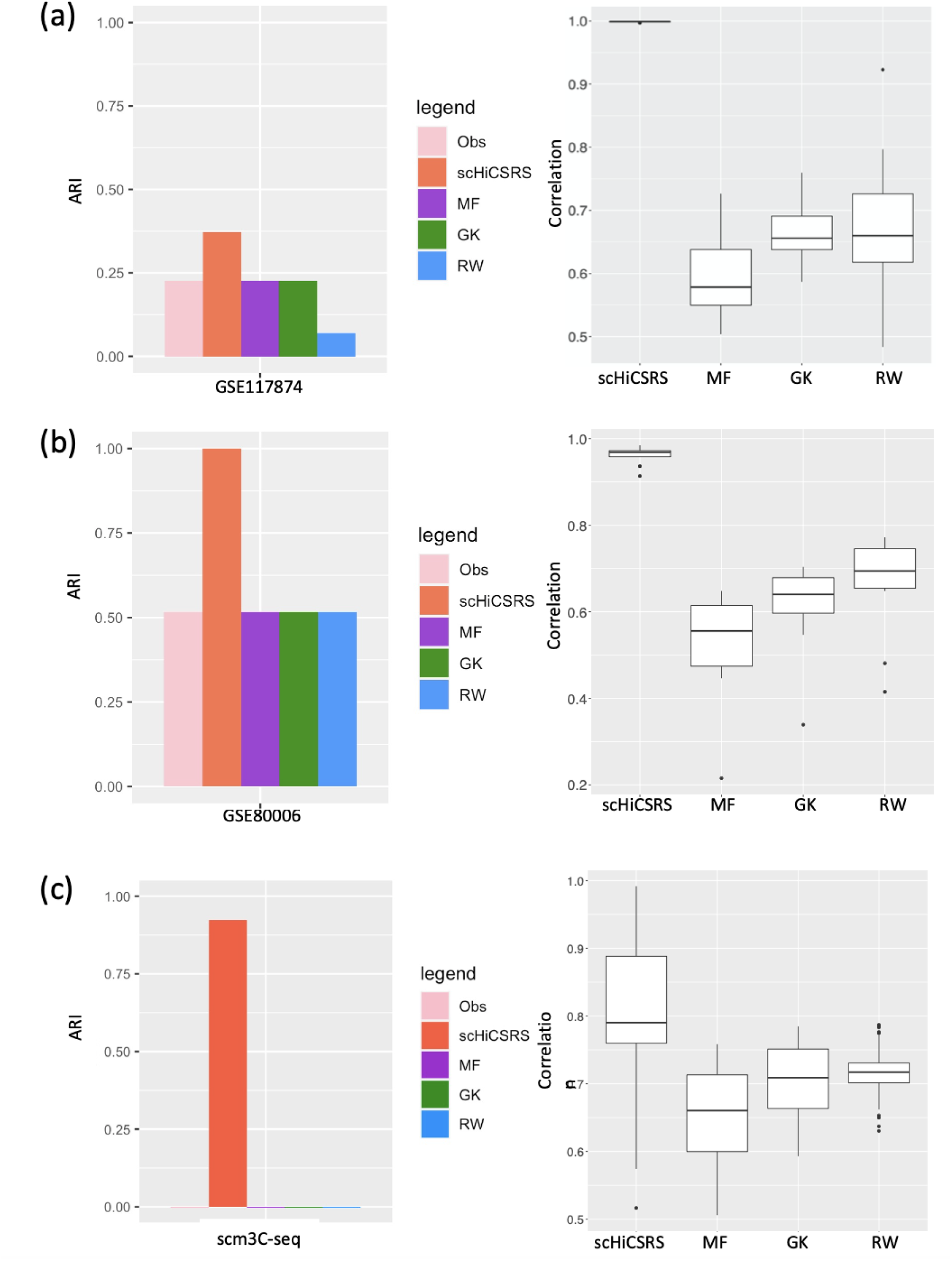
A comparison of four methods based on ARI and correlations between the imputed and nonzero observed values. The results are from the analyses of three real datasets: (a) GSE117874; (b) GSE80006; (c) scm3C-seq data.

Since structural zeros are critical for downstream analysis such as 3D structure construction (Xiao et al., 2011; Zhang et al., 2013), we are also interested in evaluating the performance of the methods when the proportion of correctly identified true structural zeros, the power, is kept at a high level. As such, we compare the performance of the four methods when the power is fixed at 0.95. For every combination of cell type, sample size, and sequencing depth, scHiCSRS maintains a much higher proportion for identifying true dropouts, the specificity (Figure 2(c) and Table S4). One can see that the overall performance of the three comparison methods, although not sensitive to the number of cells, is sensitive to the sequencing depth. In particular, for types II and III, the proportions are much smaller when the sequencing depth is 7k.

For assessing imputation accuracy, we consider the correlation between the imputed values and the expected values underlying our simulation (Figure 2(d) and Table S5). We can see that scHiCSRS has the highest correlations compared to the other methods in each of the scenarios studied. Evaluation based on the absolute error shows that it is the smallest for scHiCSRS across cell types, sample size, and sequencing depth (Figure 2(e) and Table S6), consistent with the correlation results.

## 4 Real data analysis

We consider the following three real scHi-C datasets to demonstrate the improvement of cell type clustering after data improvement with scHiCSRS and compare with the results using data improved by the three comparison methods: MF, GK, and RW.

- GSE117874: It consists of 14 GM cells (lymphoblastoid) and 18 PBMC (peripheral blood mononuclear cells) (Tan et al., 2018) (https://www.ncbi.nlm.nih.gov/geo/query/acc.cgi?acc=GSE117874). We analyzed a sub-2D matrix of dimension 30 × 30 on chromosome 1.
- GSE80006: It consists of 19 scHi-C data of K562A cells and 15 K562B cells (Flyamer et al., 2017) (https://www.ncbi.nlm.nih.gov/geo/query/acc.cgi?acc=GSE80006). Our analysis only considered the 10 single cells having a sequencing depth of at least 5K. All intra-chromosomal data from these 10 cells were used.
- scm3C-seq: It consists of scHi-C data of over 4200 single human brain prefrontal cortex cells (https://github.com/dixonlab/scm3C-seq). Eight neuronal subtypes, including L4 and L5, were all clustered together based on observed scHi-C data (Lee et al., 2019). In this analysis, we considered intra-chromosomal data of 131 cells from subtypes L4 and 180 cells from L5 that are known to be located on different cortical layers.

We explore whether the imputed data from scHiCSRS can improve downstream clustering using the K-means algorithm and assess the results based on the adjusted rand index (ARI). For GSE117874, scHiCSRS corrected two misclassifications with the original data before imputation, leading to a higher ARI (Figure **??**(a) left panel, and Table S7). MF and GK did not result in improvement, whereas RW led to more misclassifications. Further, the imputed values from scHiCSRS are much more highly correlated with the observed non-zero values across all cells (Figure **??**(a) right panel). For GSE80006, MF, GK, and RW failed to improve clustering at all while scHiCSRS corrected the misclassification, leading to an ARI of 1 and the highest correlations between the imputed and observed non-zero values (Figure **??**(b) and Table S7). The scm3C-seq dataset has many more single cells compared to the other two datasets, and the two types of cells, L4 and L5, before data improvement are highly mixed, as reflected in their near zero ARIs (Figure **??**(c) and Table S7), consistent with an earlier finding (Lee et al., 2019). Once again, scHiCSRS was able to separate most of the L4 and L5 cells, with only 6 misclassifications, leading to a much higher ARI. In contrast, there is no improvement using the enhanced data from MF, GK, and RW, where the two types of cells are still highly mixed.

## 5 Conclusion and Discussion

This paper proposes a self-representation smoothing method coupled with mixture modeling for scHi-C data quality improvement and identification of structural zeros. From both simulation and real data studies, we can see that scHiCSRS outperforms existing methods for the accuracy of imputing the contact counts of dropouts based on multiple criteria. We can also see that the Gaussian mixture model has the ability to identify structural zeros, is much better than the comparison methods using thresholding as suggested in the literature, and is not sensitive to sequencing depth. These conclusions are based on outcomes from considering several factors, including the number of cells, sequencing depth, and multiple cell types. The improved data from scHiCSRS has greatly impacted downstream analysis. From the examples of clustering GM and PBMC cells, K562 cells, and prefrontal cortex cells, we have seen that data improved with scHiCSRS led to more accurate clustering judging from known cell types.

One drawback of scHiCSRS is the large memory space it requires. As the dimension of scHi-C contact matrix increases, the memory space it requires increases exponentially, making it difficult to run on a local computer. Besides, it can be much more computationally intensive for scHiCSRS compared to the other methods, especially when the number of cells analyzed together is large, as in the case of the L4/L5 prefrontal cortex data (Table S8). This is not surprising given that for scHiCSRS, all cells are analyzed simultaneously to borrow information from one another to increase statistical power and imputation accuracy, whereas the other methods analyze each cell separately. The fuller use of available information and thus the much better performance of scHiCSRS justifies its computational cost especially since it is still practically feasible; nevertheless, effort will continue to be made to further improve computational efficiency.

## Acknowledgements

This research is supported in part by a grant from the National Institute of Health R01GM114142.

## Supplementary Document for

### Optimization procedure

We estimate the coefficient matrices *H* = {*H*_*ss’*_} and *S* = {*S*_*k’ k*_} in the self-representation smoothing model through a penalized least squared method (Jin et al., 2020). We define the following objective function:

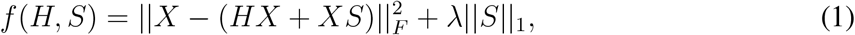

where || · ||_*F*_ and || · ||_1_ are the Frobenius and *l*_1_ norm, respectively, and *λ* is a non-negative tuning (penalty) parameter. Therefore, this may be interpreted as analogous to a Lasso type objective function. According to Gordon’s Theorem (**?**), a proper Lasso penalty parameter *λ* is at the order of the standard deviation of the noises (**?**). For simplicity and following the literature (Jin et al., 2020), we fix an estimate for *λ* before estimating the coefficient matrices. Specifically, we used *X* − *mean*(*X*) to estimate the noise matrix and set the tuning parameter as *λ* = *sd*(*X* − *mean*(*X*)) = *sd*(*X*).

A coordinate descent algorithm is used to minimize *f* (*H, S*). Specifically, we iteratively estimate one of the coefficient matrices to minimize the objective function while keeping the other one fixed. The iterative steps are as follows.

- First, we minimize equation (**??**) with respect to *S* while keeping *H* fixed:

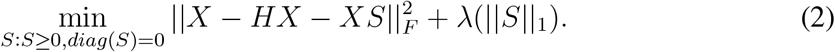
- Then we minimize equation (**??**) with respect to *H* while keeping *S* fixed, noting that the non-neighborhood positions have zero coefficients:

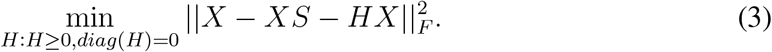

The above iterative procedure is repeated until the difference between two consecutive objective functions is less than a threshold (e.g. 0.001). The estimated data matrix, in log-normalized scale, is then 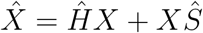.

We note that the constraints *H* ≥ 0, *S* ≥ 0 guarantee that the coefficients are non-negative and the constraints diag(*H*) = 0, diag(*S*) = 0 are used to eliminate the influence from oneself. We also note that (3) does not include a sparsity inducing term since *H* is already a sparse matrix given the typically small neighborhood constraint. Alternatively, one may set the neighborhood to be larger but include a sparsity inducing term in both equations (1) and (3).

## Supplementary Tables

**Table S1:**
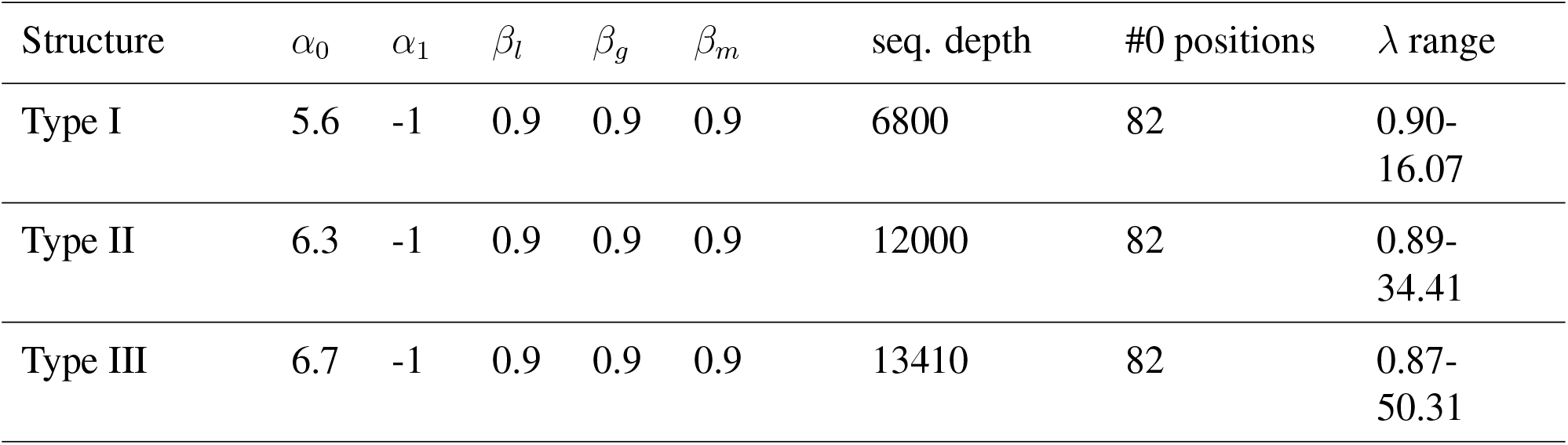
Parameter :settings for simulating scHi-C data based on three structures (Type I, II, and III) inferred from three K562 single cells.

**Table S2:**
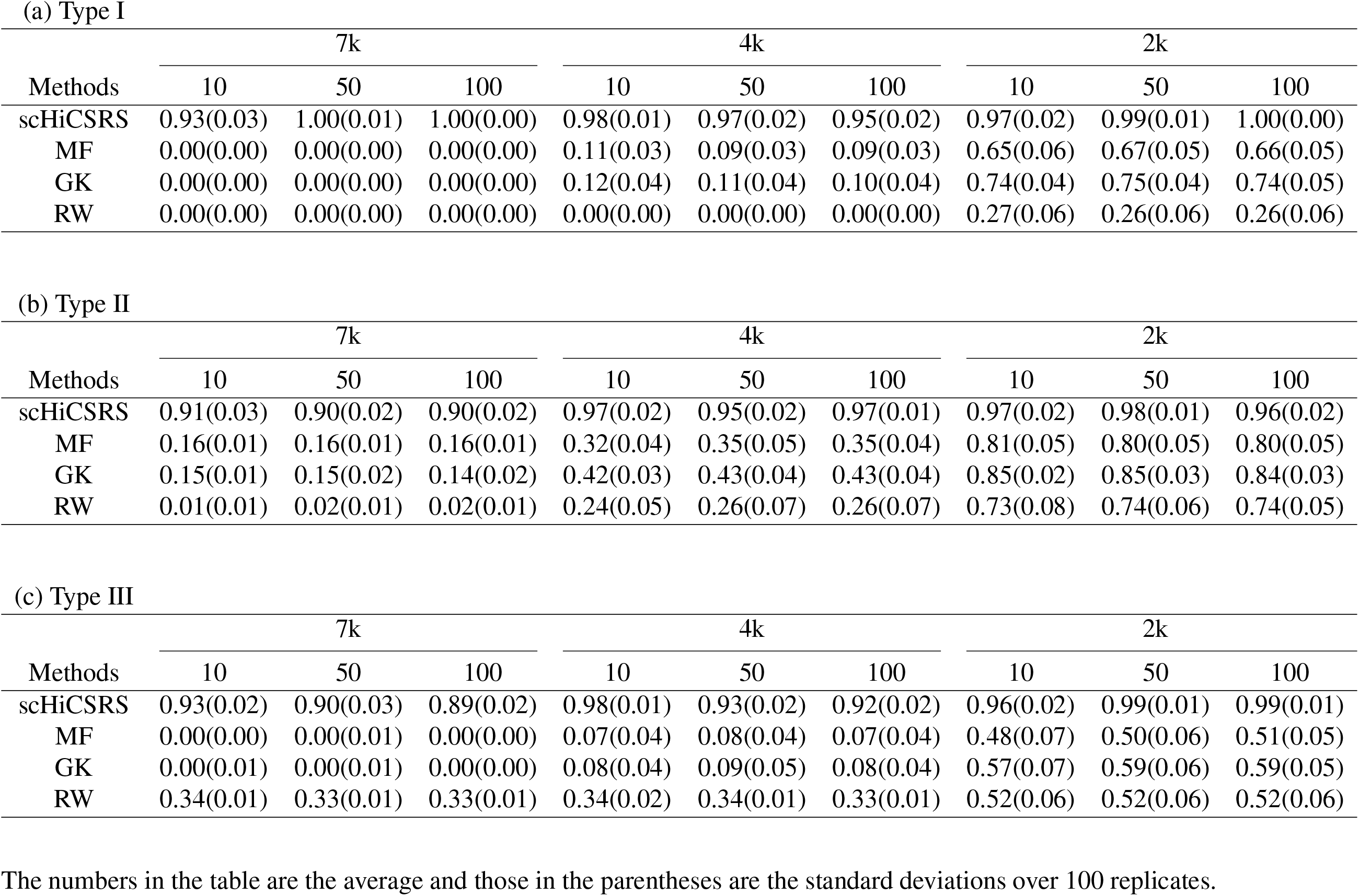
Proportion of true structural zeros correctly identified (power/sensitivity) by scHiCSRS or three comparison methods for the K562 simulated data: (a) Type I, (b) Type II, and (c) Type III.

**Table S3:**
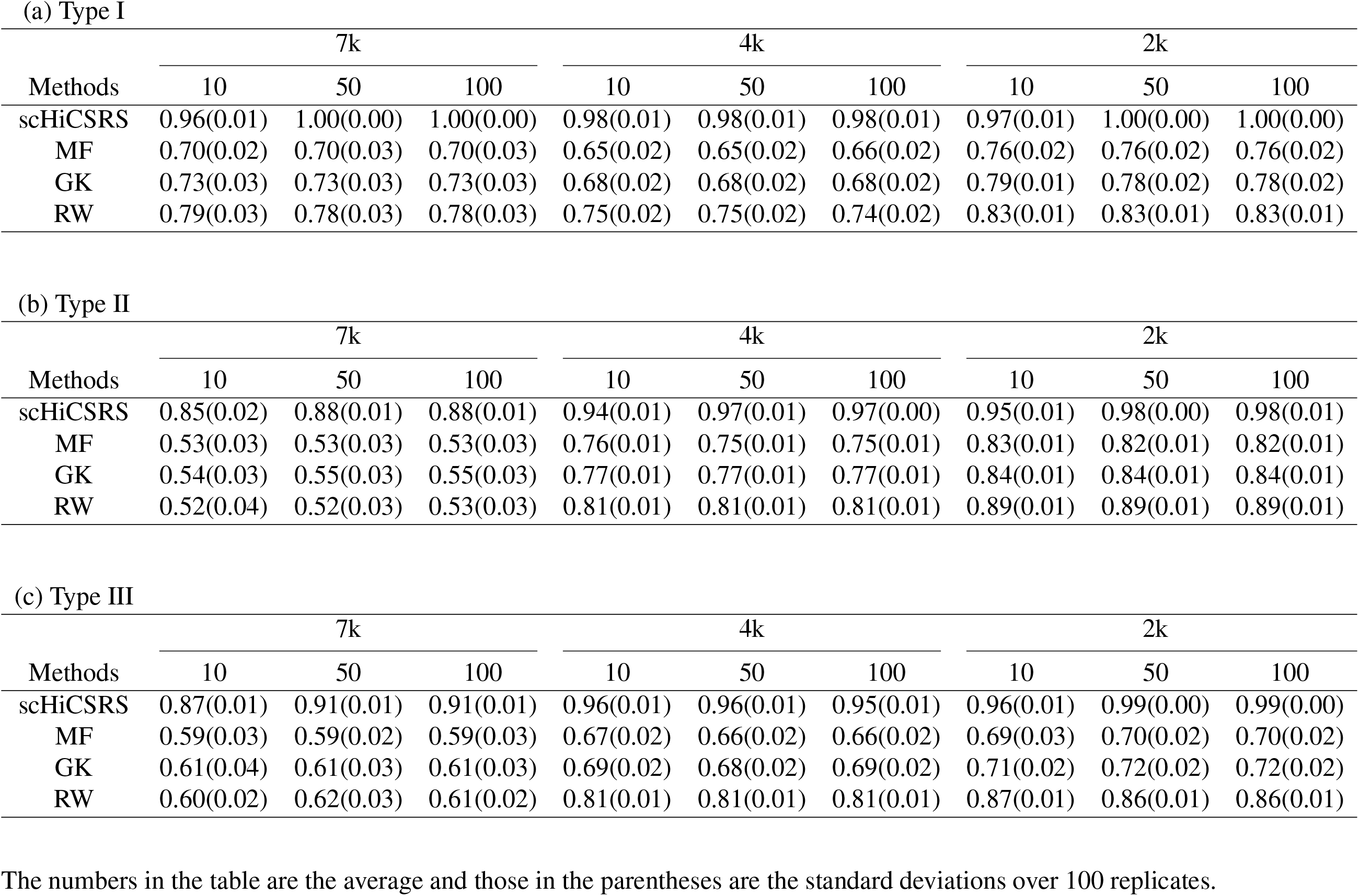
Area under the curve (AUC) criterion values for scHiCSRS and three comparison methods for the K562 simulated data:(a) Type I, (b) Type II, and (c) Type III.

**Table S4:**
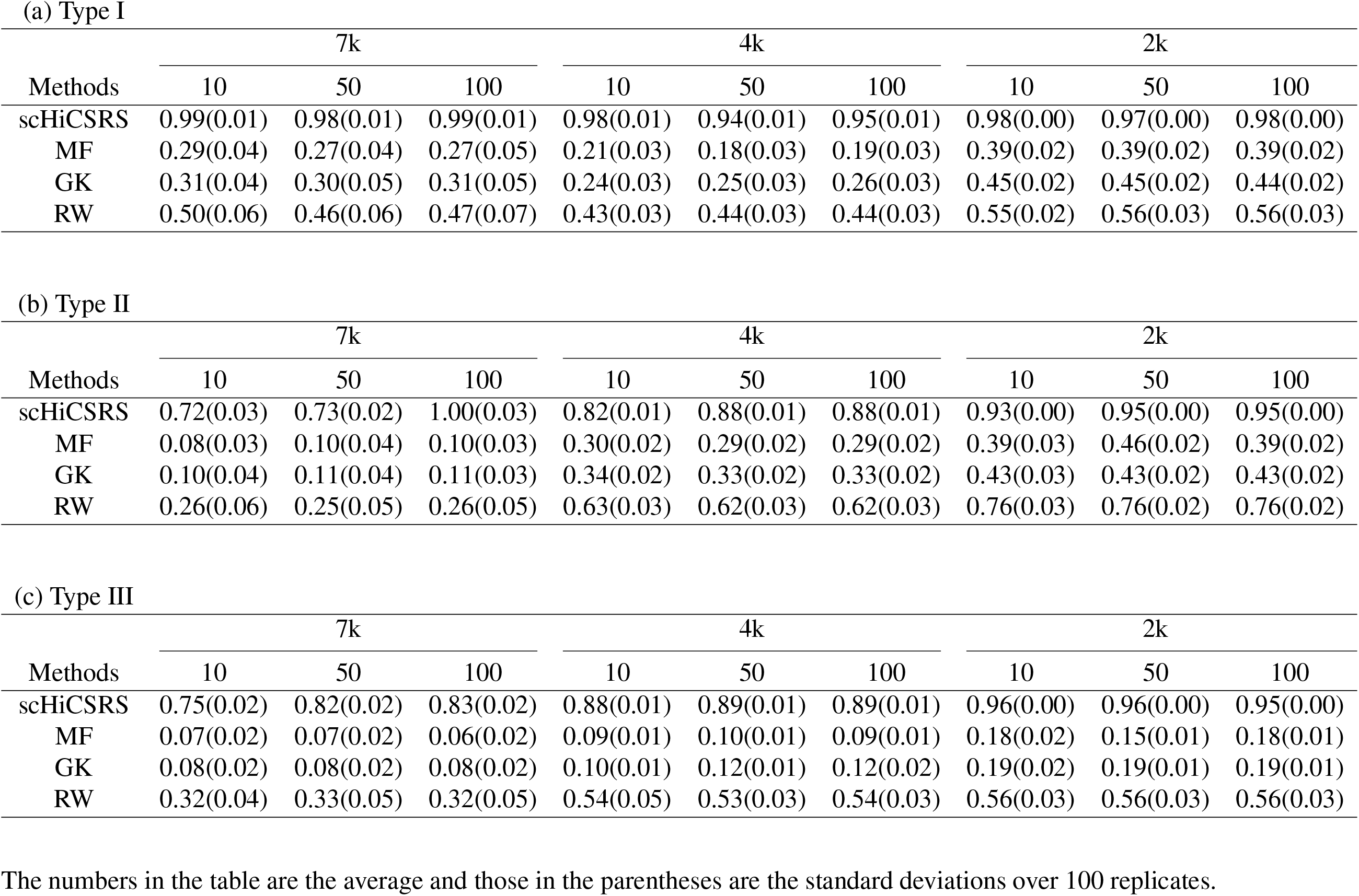
Proportion of true dropouts correctly identified (specificity) by scHiCSRS or three comparison methods for the K562 simulated data when the sensitivity is held at 0.95: (a) Type I, (b) Type II, and (c) Type III.

**Table S5:**
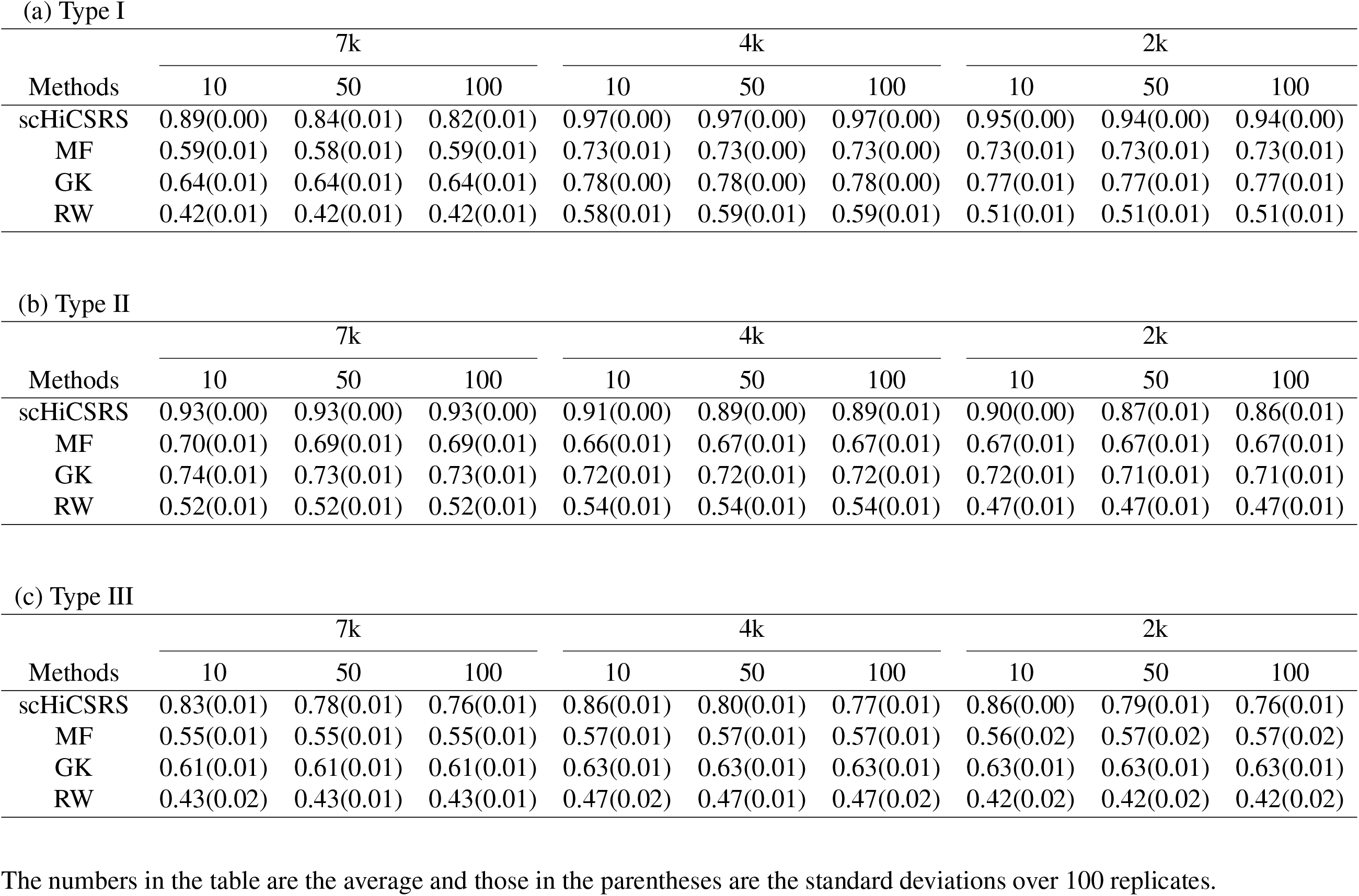
Correlation between expected and values imputed by scHiCSRS or three comparison methods for the K562 simulated data: (a) Type I, (b) Type II, and (c) Type III.

**Table S6:**
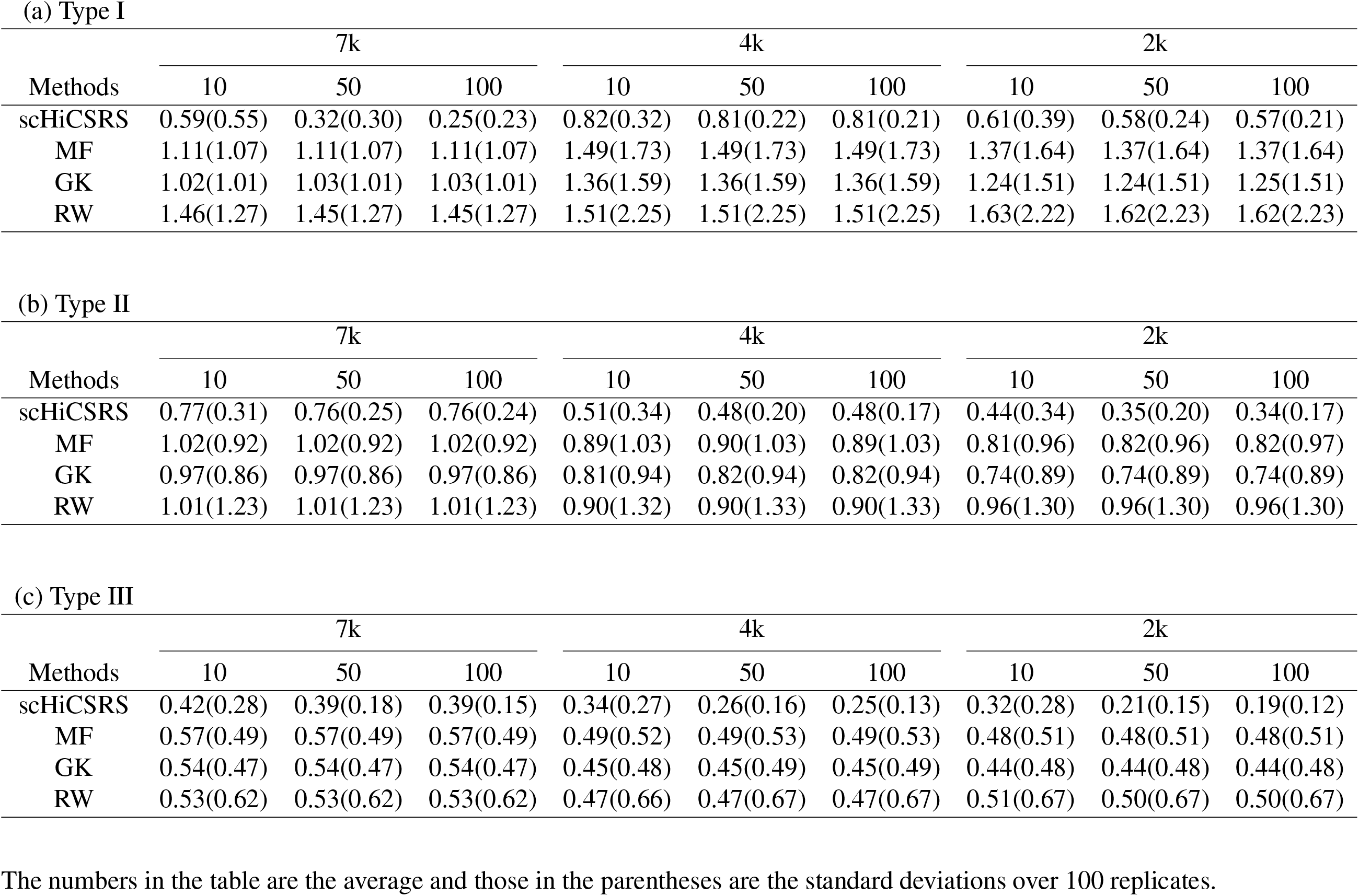
Absolute difference between the expected and the values predicted by scHiCSRS or three comparison methods for the K562 simulated data: (a) Type I, (b) Type II, and (c) Type III.

**Table S7:**
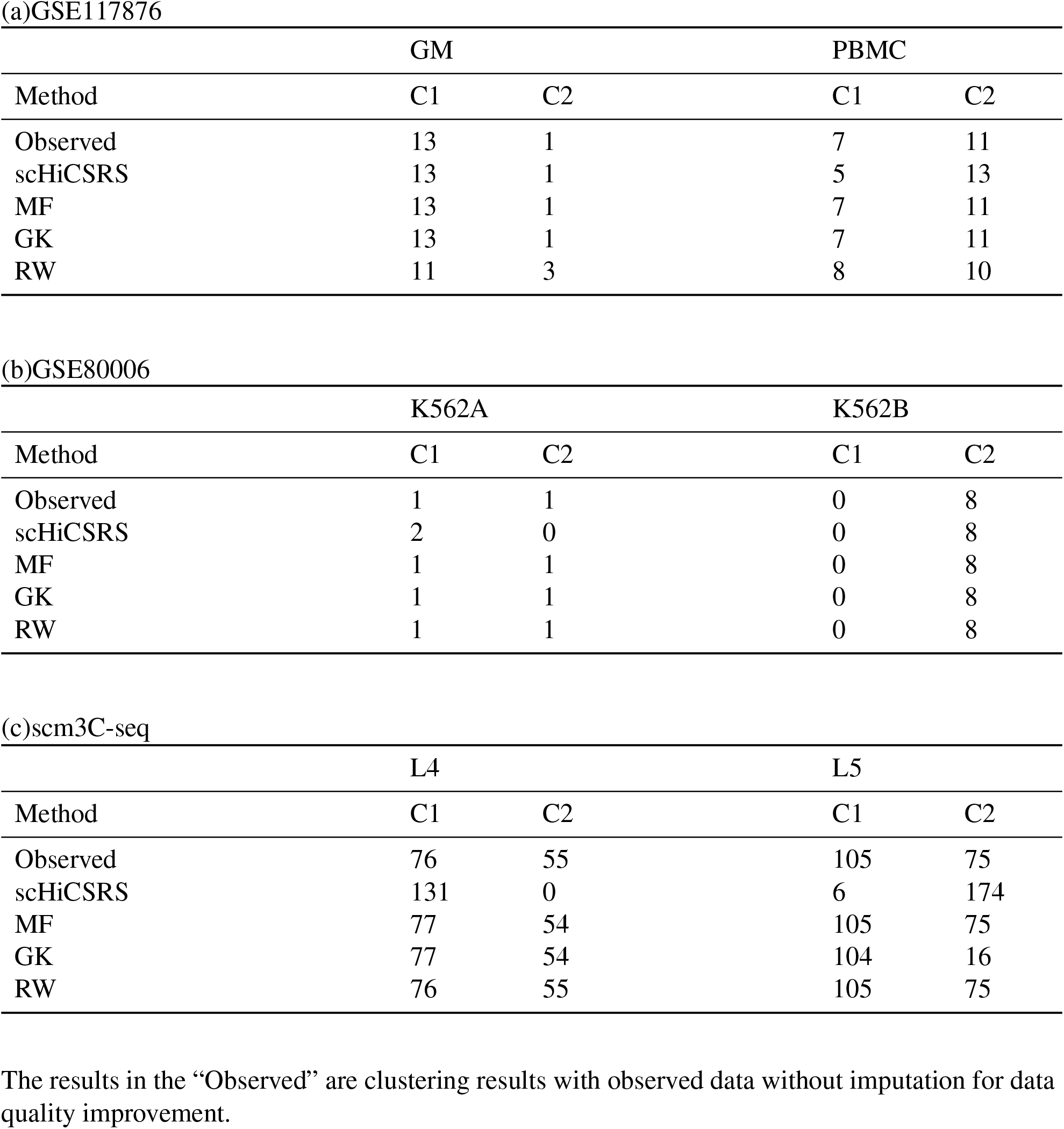
Clustering results for three single-cell Hi-C data sets.

**Table S8:**
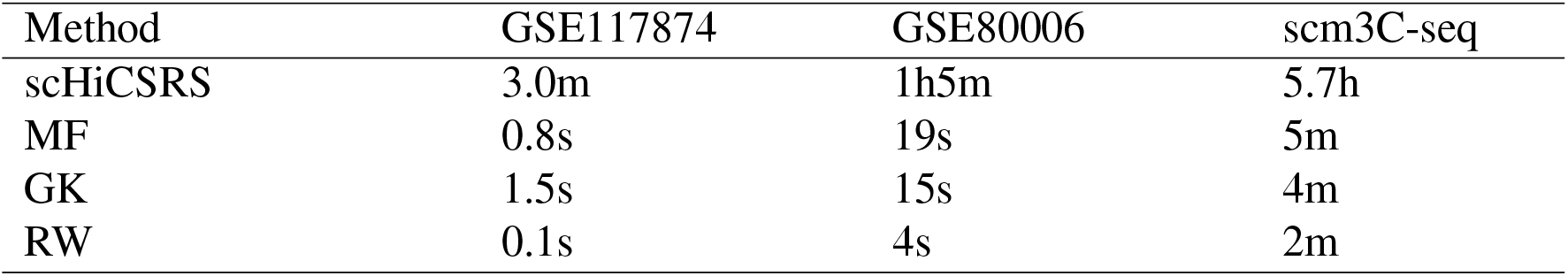
Computation time of the methods on three single cell Hi-C data sets.

## References

Job Dekker. Gene regulation in the third dimension. Science, 319(5871):1793–1794, 2008.

Erez Lieberman-Aiden, Nynke L Van Berkum, Louise Williams, Maxim Imakaev, Tobias Ragoczy, Agnes Telling, Ido Amit, Bryan R Lajoie, Peter J Sabo, Michael O Dorschner, et al. Comprehensive mapping of long-range interactions reveals folding principles of the human genome. science, 326(5950):289–293, 2009.

Suhas SP Rao, Miriam H Huntley, Neva C Durand, Elena K Stamenova, Ivan D Bochkov, James T Robinson, Adrian L Sanborn, Ido Machol, Arina D Omer, Eric S Lander, et al. A 3d map of the human genome at kilobase resolution reveals principles of chromatin looping. Cell, 159(7): 1665–1680, 2014.

Seungsoo Kim, Ivan Liachko, Donna G Brickner, Kate Cook, William S Noble, Jason H Brickner, Jay Shendure, and Maitreya J Dunham. The dynamic three-dimensional organization of the diploid yeast genome. Elife, 6:e23623, 2017.

Emily M Darrow, Miriam H Huntley, Olga Dudchenko, Elena K Stamenova, Neva C Durand, Zhuo Sun, Su-Chen Huang, Adrian L Sanborn, Ido Machol, Muhammad Shamim, et al. Deletion of dxz4 on the human inactive x chromosome alters higher-order genome architecture. Proceedings of the National Academy of Sciences, 113(31):E4504–E4512, 2016.

James Fraser, Iain Williamson, Wendy A Bickmore, and Josée Dostie. An overview of genome organization and how we got there: from fish to hi-c. Microbiol. Mol. Biol. Rev., 79(3):347–372, 2015.

Takashi Nagano, Yaniv Lubling, Tim J Stevens, Stefan Schoenfelder, Eitan Yaffe, Wendy Dean, Ernest D Laue, Amos Tanay, and Peter Fraser. Single-cell hi-c reveals cell-to-cell variability in chromosome structure. Nature, 502(7469):59–64, 2013.

Vijay Ramani, Xinxian Deng, Ruolan Qiu, Choli Lee, Christine M Disteche, William S Noble, Jay Shendure, and Zhijun Duan. Sci-hi-c: a single-cell hi-c method for mapping 3d genome organization in large number of single cells. Methods, 2019.

Takashi Nagano, Yaniv Lubling, Eitan Yaffe, Steven W Wingett, Wendy Dean, Amos Tanay, and Peter Fraser. Single-cell hi-c for genome-wide detection of chromatin interactions that occur simultaneously in a single cell. Nature protocols, 10(12):1986, 2015.

Jingtian Zhou, Jianzhu Ma, Yusi Chen, Chuankai Cheng, Bokan Bao, Jian Peng, Terrence J Sejnowski, Jesse R Dixon, and Joseph R Ecker. Robust single-cell hi-c clustering by convolution- and random-walk–based imputation. Proceedings of the National Academy of Sciences, page 201901423, 2019.

David van Dijk, Juozas Nainys, Roshan Sharma, Pooja Kathail, Ambrose J Carr, Kevin R Moon, Linas Mazutis, Guy Wolf, Smita Krishnaswamy, and Dana Pe’er. Magic: A diffusion-based imputation method reveals gene-gene interactions in single-cell rna-sequencing data. BioRxiv, page 111591, 2017.

Chong Chen, Changjing Wu, Linjie Wu, Yishu Wang, Minghua Deng, and Ruibin Xi. scrmd: Imputation for single cell rna-seq data via robust matrix decomposition. bioRxiv, page 459404, 2018.

Wei Vivian Li and Jingyi Jessica Li. An accurate and robust imputation method scimpute for single-cell rna-seq data. Nature communications, 9(1):1–9, 2018.

Aanchal Mongia, Debarka Sengupta, and Angshul Majumdar. Mcimpute: Matrix completion based imputation for single cell rna-seq data. Frontiers in genetics, 10:9, 2019.

Tao Peng, Qin Zhu, Penghang Yin, and Kai Tan. Scrabble: single-cell rna-seq imputation constrained by bulk rna-seq data. Genome biology, 20(1):88, 2019.

Yinlei Hu, Bin Li, Wen Zhang, Nianping Liu, Pengfei Cai, Falai Chen, and Kun Qu. Wedge: imputation of gene expression values from single-cell rna-seq datasets using biased matrix decomposition. bioRxiv, page 864488, 2020.

Xiang Zhou, Hua Chai, Huiying Zhao, Ching-Hsing Luo, and Yuedong Yang. Imputing missing rna-sequencing data from dna methylation by using a transfer learning–based neural network. GigaScience, 9(7):giaa076, 2020.

Maryam Zand and Jianhua Ruan. Network-based single-cell rna-seq data imputation enhances cell type identification. Genes, 11(4):377, 2020.

Jiahua Rao, Xiang Zhou, Yutong Lu, Huiying Zhao, and Yuedong Yang. Imputing single-cell rna-seq data by combining graph convolution and autoencoder neural networks. iScience, page 102393, 2021.

Tao Yang, Feipeng Zhang, Galip Gürkan Yardımcı, Fan Song, Ross C Hardison, William Stafford Noble, Feng Yue, and Qunhua Li. Hicrep: assessing the reproducibility of hi-c data using a stratum-adjusted correlation coefficient. Genome research, 27(11):1939–1949, 2017.

Oana Ursu, Nathan Boley, Maryna Taranova, YX Rachel Wang, Galip Gurkan Yardimci, William Stafford Noble, and Anshul Kundaje. Genomedisco: A concordance score for chromosome conformation capture experiments using random walks on contact map graphs. Bioinformatics, 34(16):2701–2707, 2018.

Hao Zhu and Zheng Wang. Scl: a lattice-based approach to infer 3d chromosome structures from single-cell hi-c data. Bioinformatics, 35(20):3981–3988, 2019.

Chenggong Han, Qing Xie, and Shili Lin. Are dropout imputation methods for scrna-seq effective for schi-c data? Briefings in Bioinformatics, 2020.

Caiwei Zhen, Yuxian Wang, Lu Han, Jingyi Li, Jinghao Peng, Tao Wang, Jianye Hao, Xuequn Shang, Zhongyu Wei, and Jiajie Peng. A novel framework for single-cell hi-c clustering based on graph-convolution-based imputation and two-phase-based feature extraction. bioRxiv, 2021.

Yan Zhang, Lin An, Jie Xu, Bo Zhang, W Jim Zheng, Ming Hu, Jijun Tang, and Feng Yue. Enhancing hi-c data resolution with deep convolutional neural network hicplus. Nature communications, 9(1):750, 2018.

Hao Hong, Shuai Jiang, Hao Li, Guifang Du, Yu Sun, Huan Tao, Cheng Quan, Chenghui Zhao, Ruijiang Li, Wanying Li, et al. Deephic: A generative adversarial network for enhancing hi-c data resolution. PLoS computational biology, 16(2):e1007287, 2020.

Ke Jin, Le Ou-Yang, Xing-Ming Zhao, Hong Yan, and Xiao-Fei Zhang. sctssr: gene expression recovery for single-cell rna sequencing using two-side sparse self-representation. Bioinformatics, 36(10):3131–3138, 2020.

Mo Huang, Jingshu Wang, Eduardo Torre, Hannah Dueck, Sydney Shaffer, Roberto Bonasio, John Murray, Arjun Raj, Mingyao Li, and Nancy R Zhang. Gene expression recovery for single cell rna sequencing. bioRxiv, page 138677, 2017.

Gerda Claeskens, Nils Lid Hjort, et al. Model selection and model averaging. Cambridge Books, 2008.

Michael Rosenthal, Darshan Bryner, Fred Huffer, Shane Evans, Anuj Srivastava, and Nicola Neretti. Bayesian estimation of three-dimensional chromosomal structure from single-cell hi-c data. Journal of Computational Biology, 2019.

Ilya M Flyamer, Johanna Gassler, Maxim Imakaev, Hugo B Brandão, Sergey V Ulianov, Nezar Abdennur, Sergey V Razin, Leonid A Mirny, and Kikuë Tachibana-Konwalski. Single-nucleus hi-c reveals unique chromatin reorganization at oocyte-to-zygote transition. Nature, 544(7648): 110–114, 2017.

Jincheol Park and Shili Lin. Evaluation and comparison of methods for recapitulation of 3d spatial chromatin structures. Briefings in bioinformatics, 20(4):1205–1214, 2019.

Guanghua Xiao, Xinlei Wang, and Arkady B Khodursky. Modeling three-dimensional chromosome structures using gene expression data. Journal of the American Statistical Association, 106(493): 61–72, 2011.

ZhiZhuo Zhang, Guoliang Li, Kim-Chuan Toh, and Wing-Kin Sung. Inference of spatial organizations of chromosomes using semi-definite embedding approach and hi-c data. In Annual international conference on research in computational molecular biology, pages 317–332. Springer, 2013.

Longzhi Tan, Dong Xing, Chi-Han Chang, Heng Li, and X Sunney Xie. Three-dimensional genome structures of single diploid human cells. Science, 361(6405):924–928, 2018.

Dong-Sung Lee, Chongyuan Luo, Jingtian Zhou, Sahaana Chandran, Angeline Rivkin, Anna Bartlett, Joseph R Nery, Conor Fitzpatrick, Carolyn O’Connor, Jesse R Dixon, et al. Simul-taneous profiling of 3d genome structure and dna methylation in single human cells. Nature methods, 16(10):999–1006, 2019.

