## Supplementary for "scHiCSRS: A Self-Representation Smoothing Method with Gaussian Mixture Model for Imputing single cell Hi-C Data"

### Optimization procedure

We estimate the coefficient matrices $H = \{H_{ss'}\}$ and $S = \{S_{k'k}\}$ in the self-representation smoothing model through a penalized least squared method (Jin et al., 2020). We define the following objective function:

$$f(H, S) = ||X - (HX + XS)||_F^2 + \lambda||S||_1, \tag{1}$$

where $||\cdot||_F$ and $||\cdot||_1$ are the Frobenius and $l_1$ norm, respectively, and $\lambda$ is a non-negative tuning (penalty) parameter. Therefore, this may be interpreted as analogous to a Lasso type objective function. According to Gordon's Theorem (Vershynin, 2010), a proper Lasso penalty parameter $\lambda$ is at the order of the standard deviation of the noises (Zhao et al., 2017). For simplicity and following the literature (Jin et al., 2020), we fix an estimate for $\lambda$ before estimating the coefficient matrices. Specifically, we used $X - mean(X)$ to estimate the noise matrix and set the tuning parameter as $\lambda = sd(X - mean(X)) = sd(X)$.

A coordinate descent algorithm is used to minimize $f(H, S)$. Specifically, we iteratively estimate one of the coefficient matrices to minimize the objective function while keeping the other one fixed. The iterative steps are as follows.

- First, we minimize equation (1) with respect to $S$ while keeping $H$ fixed:

$$\min_{S:S\geq 0, diag(S)=0} ||X - HX - XS||_F^2 + \lambda(||S||_1). \tag{2}$$

- Then we minimize equation (1) with respect to $H$ while keeping $S$ fixed, noting that the non-neighborhood positions have zero coefficients:

$$\min_{H:H\geq 0, diag(H)=0} ||X - XS - HX||_F^2. \tag{3}$$

The above iterative procedure is repeated until the difference between two consecutive objective functions is less than a threshold (e.g. 0.001). The estimated data matrix, in log-normalized scale, is then $\hat{X} = \hat{H}X + X\hat{S}$.

We note that the constraints $H \geq 0, S \geq 0$ guarantee that the coefficients are non-negative and the constraints $\text{diag}(H) = 0$, $\text{diag}(S) = 0$ are used to eliminate the influence from oneself. We also note that (3) does not include a sparsity inducing term since $H$ is already a sparse matrix given the typically small neighborhood constraint. Alternatively, one may set the neighborhood to be larger but include a sparsity inducing term in both equations (1) and (3).

### Supplementary Tables

Table S1: Parameter :settings for simulating scHi-C data based on three structures (Type I, II, and III) inferred from three K562 single cells.

| Structure | $\alpha_0$ | $\alpha_1$ | $\beta_l$ | $\beta_g$ | $\beta_m$ | seq. depth | #0 positions | $\lambda$ range |
|---|---|---|---|---|---|---|---|---|
| Type I | 5.6 | -1 | 0.9 | 0.9 | 0.9 | 6800 | 82 | 0.90-16.07 |
| Type II | 6.3 | -1 | 0.9 | 0.9 | 0.9 | 12000 | 82 | 0.89-34.41 |
| Type III | 6.7 | -1 | 0.9 | 0.9 | 0.9 | 13410 | 82 | 0.87-50.31 |

Table S2: Proportion of true structural zeros correctly identified (power/sensitivity) by scHiCSRS or three comparison methods for the K562 simulated data: (a) Type I, (b) Type II, and (c) Type III.

(a) Type I

| | 7k | | | 4k | | | 2k | | |
|---|---|---|---|---|---|---|---|---|---|
| Methods | 10 | 50 | 100 | 10 | 50 | 100 | 10 | 50 | 100 |
| scHiCSRS | 0.93(0.03) | 1.00(0.01) | 1.00(0.00) | 0.98(0.01) | 0.97(0.02) | 0.95(0.02) | 0.97(0.02) | 0.99(0.01) | 1.00(0.00) |
| MF | 0.00(0.00) | 0.00(0.00) | 0.00(0.00) | 0.11(0.03) | 0.09(0.03) | 0.09(0.03) | 0.65(0.06) | 0.67(0.05) | 0.66(0.05) |
| GK | 0.00(0.00) | 0.00(0.00) | 0.00(0.00) | 0.12(0.04) | 0.11(0.04) | 0.10(0.04) | 0.74(0.04) | 0.75(0.04) | 0.74(0.05) |
| RW | 0.00(0.00) | 0.00(0.00) | 0.00(0.00) | 0.00(0.00) | 0.00(0.00) | 0.00(0.00) | 0.27(0.06) | 0.26(0.06) | 0.26(0.06) |

(b) Type II

| | 7k | | | 4k | | | 2k | | |
|---|---|---|---|---|---|---|---|---|---|
| Methods | 10 | 50 | 100 | 10 | 50 | 100 | 10 | 50 | 100 |
| scHiCSRS | 0.91(0.03) | 0.90(0.02) | 0.90(0.02) | 0.97(0.02) | 0.95(0.02) | 0.97(0.01) | 0.97(0.02) | 0.98(0.01) | 0.96(0.02) |
| MF | 0.16(0.01) | 0.16(0.01) | 0.16(0.01) | 0.32(0.04) | 0.35(0.05) | 0.35(0.04) | 0.81(0.05) | 0.80(0.05) | 0.80(0.05) |
| GK | 0.15(0.01) | 0.15(0.02) | 0.14(0.02) | 0.42(0.03) | 0.43(0.04) | 0.43(0.04) | 0.85(0.02) | 0.85(0.03) | 0.84(0.03) |
| RW | 0.01(0.01) | 0.02(0.01) | 0.02(0.01) | 0.24(0.05) | 0.26(0.07) | 0.26(0.07) | 0.73(0.08) | 0.74(0.06) | 0.74(0.05) |

(c) Type III

| | 7k | | | 4k | | | 2k | | |
|---|---|---|---|---|---|---|---|---|---|
| Methods | 10 | 50 | 100 | 10 | 50 | 100 | 10 | 50 | 100 |
| scHiCSRS | 0.93(0.02) | 0.90(0.03) | 0.89(0.02) | 0.98(0.01) | 0.93(0.02) | 0.92(0.02) | 0.96(0.02) | 0.99(0.01) | 0.99(0.01) |
| MF | 0.00(0.00) | 0.00(0.01) | 0.00(0.00) | 0.07(0.04) | 0.08(0.04) | 0.07(0.04) | 0.48(0.07) | 0.50(0.06) | 0.51(0.05) |
| GK | 0.00(0.01) | 0.00(0.01) | 0.00(0.00) | 0.08(0.04) | 0.09(0.05) | 0.08(0.04) | 0.57(0.07) | 0.59(0.06) | 0.59(0.05) |
| RW | 0.34(0.01) | 0.33(0.01) | 0.33(0.01) | 0.34(0.02) | 0.34(0.01) | 0.33(0.01) | 0.52(0.06) | 0.52(0.06) | 0.52(0.06) |

The numbers in the table are the average and those in the parentheses are the standard deviations over 100 replicates.

Table S3: Area under the curve (AUC) criterion values for scHiCSRS and three comparison methods for the K562 simulated data: (a) Type I, (b) Type II, and (c) Type III.

(a) Type I

| | 7k | | | 4k | | | 2k | | |
|---|---|---|---|---|---|---|---|---|---|
| Methods | 10 | 50 | 100 | 10 | 50 | 100 | 10 | 50 | 100 |
| scHiCSRS | 0.96(0.01) | 1.00(0.00) | 1.00(0.00) | 0.98(0.01) | 0.98(0.01) | 0.98(0.01) | 0.97(0.01) | 1.00(0.00) | 1.00(0.00) |
| MF | 0.70(0.02) | 0.70(0.03) | 0.70(0.03) | 0.65(0.02) | 0.65(0.02) | 0.66(0.02) | 0.76(0.02) | 0.76(0.02) | 0.76(0.02) |
| GK | 0.73(0.03) | 0.73(0.03) | 0.73(0.03) | 0.68(0.02) | 0.68(0.02) | 0.68(0.02) | 0.79(0.01) | 0.78(0.02) | 0.78(0.02) |
| RW | 0.79(0.03) | 0.78(0.03) | 0.78(0.03) | 0.75(0.02) | 0.75(0.02) | 0.74(0.02) | 0.83(0.01) | 0.83(0.01) | 0.83(0.01) |

(b) Type II

| | 7k | | | 4k | | | 2k | | |
|---|---|---|---|---|---|---|---|---|---|
| Methods | 10 | 50 | 100 | 10 | 50 | 100 | 10 | 50 | 100 |
| scHiCSRS | 0.85(0.02) | 0.88(0.01) | 0.88(0.01) | 0.94(0.01) | 0.97(0.01) | 0.97(0.00) | 0.95(0.01) | 0.98(0.00) | 0.98(0.01) |
| MF | 0.53(0.03) | 0.53(0.03) | 0.53(0.03) | 0.76(0.01) | 0.75(0.01) | 0.75(0.01) | 0.83(0.01) | 0.82(0.01) | 0.82(0.01) |
| GK | 0.54(0.03) | 0.55(0.03) | 0.55(0.03) | 0.77(0.01) | 0.77(0.01) | 0.77(0.01) | 0.84(0.01) | 0.84(0.01) | 0.84(0.01) |
| RW | 0.52(0.04) | 0.52(0.03) | 0.53(0.03) | 0.81(0.01) | 0.81(0.01) | 0.81(0.01) | 0.89(0.01) | 0.89(0.01) | 0.89(0.01) |

(c) Type III

| | 7k | | | 4k | | | 2k | | |
|---|---|---|---|---|---|---|---|---|---|
| Methods | 10 | 50 | 100 | 10 | 50 | 100 | 10 | 50 | 100 |
| scHiCSRS | 0.87(0.01) | 0.91(0.01) | 0.91(0.01) | 0.96(0.01) | 0.96(0.01) | 0.95(0.01) | 0.96(0.01) | 0.99(0.00) | 0.99(0.00) |
| MF | 0.59(0.03) | 0.59(0.02) | 0.59(0.03) | 0.67(0.02) | 0.66(0.02) | 0.66(0.02) | 0.69(0.03) | 0.70(0.02) | 0.70(0.02) |
| GK | 0.61(0.04) | 0.61(0.03) | 0.61(0.03) | 0.69(0.02) | 0.68(0.02) | 0.69(0.02) | 0.71(0.02) | 0.72(0.02) | 0.72(0.02) |
| RW | 0.60(0.02) | 0.62(0.03) | 0.61(0.02) | 0.81(0.01) | 0.81(0.01) | 0.81(0.01) | 0.87(0.01) | 0.86(0.01) | 0.86(0.01) |

The numbers in the table are the average and those in the parentheses are the standard deviations over 100 replicates.

Table S4: Proportion of true dropouts correctly identified (specificity) by scHiCSRS or three comparison methods for the K562 simulated data when the sensitivity is held at 0.95: (a) Type I, (b) Type II, and (c) Type III.

(a) Type I

| | 7k | | | 4k | | | 2k | | |
|---|---|---|---|---|---|---|---|---|---|
| Methods | 10 | 50 | 100 | 10 | 50 | 100 | 10 | 50 | 100 |
| scHiCSRS | 0.99(0.01) | 0.98(0.01) | 0.99(0.01) | 0.98(0.01) | 0.94(0.01) | 0.95(0.01) | 0.98(0.00) | 0.97(0.00) | 0.98(0.00) |
| MF | 0.29(0.04) | 0.27(0.04) | 0.27(0.05) | 0.21(0.03) | 0.18(0.03) | 0.19(0.03) | 0.39(0.02) | 0.39(0.02) | 0.39(0.02) |
| GK | 0.31(0.04) | 0.30(0.05) | 0.31(0.05) | 0.24(0.03) | 0.25(0.03) | 0.26(0.03) | 0.45(0.02) | 0.45(0.02) | 0.44(0.02) |
| RW | 0.50(0.06) | 0.46(0.06) | 0.47(0.07) | 0.43(0.03) | 0.44(0.03) | 0.44(0.03) | 0.55(0.02) | 0.56(0.03) | 0.56(0.03) |

(b) Type II

| | 7k | | | 4k | | | 2k | | |
|---|---|---|---|---|---|---|---|---|---|
| Methods | 10 | 50 | 100 | 10 | 50 | 100 | 10 | 50 | 100 |
| scHiCSRS | 0.72(0.03) | 0.73(0.02) | 1.00(0.03) | 0.82(0.01) | 0.88(0.01) | 0.88(0.01) | 0.93(0.00) | 0.95(0.00) | 0.95(0.00) |
| MF | 0.08(0.03) | 0.10(0.04) | 0.10(0.03) | 0.30(0.02) | 0.29(0.02) | 0.29(0.02) | 0.39(0.03) | 0.46(0.02) | 0.39(0.02) |
| GK | 0.10(0.04) | 0.11(0.04) | 0.11(0.03) | 0.34(0.02) | 0.33(0.02) | 0.33(0.02) | 0.43(0.03) | 0.43(0.02) | 0.43(0.02) |
| RW | 0.26(0.06) | 0.25(0.05) | 0.26(0.05) | 0.63(0.03) | 0.62(0.03) | 0.62(0.03) | 0.76(0.03) | 0.76(0.02) | 0.76(0.02) |

(c) Type III

| | 7k | | | 4k | | | 2k | | |
|---|---|---|---|---|---|---|---|---|---|
| Methods | 10 | 50 | 100 | 10 | 50 | 100 | 10 | 50 | 100 |
| scHiCSRS | 0.75(0.02) | 0.82(0.02) | 0.83(0.02) | 0.88(0.01) | 0.89(0.01) | 0.89(0.01) | 0.96(0.00) | 0.96(0.00) | 0.95(0.00) |
| MF | 0.07(0.02) | 0.07(0.02) | 0.06(0.02) | 0.09(0.01) | 0.10(0.01) | 0.09(0.01) | 0.18(0.02) | 0.15(0.01) | 0.18(0.01) |
| GK | 0.08(0.02) | 0.08(0.02) | 0.08(0.02) | 0.10(0.01) | 0.12(0.01) | 0.12(0.02) | 0.19(0.02) | 0.19(0.01) | 0.19(0.01) |
| RW | 0.32(0.04) | 0.33(0.05) | 0.32(0.05) | 0.54(0.05) | 0.53(0.03) | 0.54(0.03) | 0.56(0.03) | 0.56(0.03) | 0.56(0.03) |

The numbers in the table are the average and those in the parentheses are the standard deviations over 100 replicates.

Table S5: Correlation between expected and values imputed by scHiCSRS or three comparison methods for the K562 simulated data: (a) Type I, (b) Type II, and (c) Type III.

(a) Type I

| | 7k | | | 4k | | | 2k | | |
|---|---|---|---|---|---|---|---|---|---|
| Methods | 10 | 50 | 100 | 10 | 50 | 100 | 10 | 50 | 100 |
| scHiCSRS | 0.89(0.00) | 0.84(0.01) | 0.82(0.01) | 0.97(0.00) | 0.97(0.00) | 0.97(0.00) | 0.95(0.00) | 0.94(0.00) | 0.94(0.00) |
| MF | 0.59(0.01) | 0.58(0.01) | 0.59(0.01) | 0.73(0.01) | 0.73(0.00) | 0.73(0.00) | 0.73(0.01) | 0.73(0.01) | 0.73(0.01) |
| GK | 0.64(0.01) | 0.64(0.01) | 0.64(0.01) | 0.78(0.00) | 0.78(0.00) | 0.78(0.00) | 0.77(0.01) | 0.77(0.01) | 0.77(0.01) |
| RW | 0.42(0.01) | 0.42(0.01) | 0.42(0.01) | 0.58(0.01) | 0.59(0.01) | 0.59(0.01) | 0.51(0.01) | 0.51(0.01) | 0.51(0.01) |

(b) Type II

| | 7k | | | 4k | | | 2k | | |
|---|---|---|---|---|---|---|---|---|---|
| Methods | 10 | 50 | 100 | 10 | 50 | 100 | 10 | 50 | 100 |
| scHiCSRS | 0.93(0.00) | 0.93(0.00) | 0.93(0.00) | 0.91(0.00) | 0.89(0.00) | 0.89(0.01) | 0.90(0.00) | 0.87(0.01) | 0.86(0.01) |
| MF | 0.70(0.01) | 0.69(0.01) | 0.69(0.01) | 0.66(0.01) | 0.67(0.01) | 0.67(0.01) | 0.67(0.01) | 0.67(0.01) | 0.67(0.01) |
| GK | 0.74(0.01) | 0.73(0.01) | 0.73(0.01) | 0.72(0.01) | 0.72(0.01) | 0.72(0.01) | 0.72(0.01) | 0.71(0.01) | 0.71(0.01) |
| RW | 0.52(0.01) | 0.52(0.01) | 0.52(0.01) | 0.54(0.01) | 0.54(0.01) | 0.54(0.01) | 0.47(0.01) | 0.47(0.01) | 0.47(0.01) |

(c) Type III

| | 7k | | | 4k | | | 2k | | |
|---|---|---|---|---|---|---|---|---|---|
| Methods | 10 | 50 | 100 | 10 | 50 | 100 | 10 | 50 | 100 |
| scHiCSRS | 0.83(0.01) | 0.78(0.01) | 0.76(0.01) | 0.86(0.01) | 0.80(0.01) | 0.77(0.01) | 0.86(0.00) | 0.79(0.01) | 0.76(0.01) |
| MF | 0.55(0.01) | 0.55(0.01) | 0.55(0.01) | 0.57(0.01) | 0.57(0.01) | 0.57(0.01) | 0.56(0.02) | 0.57(0.02) | 0.57(0.02) |
| GK | 0.61(0.01) | 0.61(0.01) | 0.61(0.01) | 0.63(0.01) | 0.63(0.01) | 0.63(0.01) | 0.63(0.01) | 0.63(0.01) | 0.63(0.01) |
| RW | 0.43(0.02) | 0.43(0.01) | 0.43(0.01) | 0.47(0.02) | 0.47(0.01) | 0.47(0.02) | 0.42(0.02) | 0.42(0.02) | 0.42(0.02) |

The numbers in the table are the average and those in the parentheses are the standard deviations over 100 replicates.

Table S6: Absolute difference between the expected and the values predicted by scHiCSRS or three comparison methods for the K562 simulated data: (a) Type I, (b) Type II, and (c) Type III.

(a) Type I

| | 7k | | | 4k | | | 2k | | |
|---|---|---|---|---|---|---|---|---|---|
| Methods | 10 | 50 | 100 | 10 | 50 | 100 | 10 | 50 | 100 |
| scHiCSRS | 0.59(0.55) | 0.32(0.30) | 0.25(0.23) | 0.82(0.32) | 0.81(0.22) | 0.81(0.21) | 0.61(0.39) | 0.58(0.24) | 0.57(0.21) |
| MF | 1.11(1.07) | 1.11(1.07) | 1.11(1.07) | 1.49(1.73) | 1.49(1.73) | 1.49(1.73) | 1.37(1.64) | 1.37(1.64) | 1.37(1.64) |
| GK | 1.02(1.01) | 1.03(1.01) | 1.03(1.01) | 1.36(1.59) | 1.36(1.59) | 1.36(1.59) | 1.24(1.51) | 1.24(1.51) | 1.25(1.51) |
| RW | 1.46(1.27) | 1.45(1.27) | 1.45(1.27) | 1.51(2.25) | 1.51(2.25) | 1.51(2.25) | 1.63(2.22) | 1.62(2.23) | 1.62(2.23) |

(b) Type II

| | 7k | | | 4k | | | 2k | | |
|---|---|---|---|---|---|---|---|---|---|
| Methods | 10 | 50 | 100 | 10 | 50 | 100 | 10 | 50 | 100 |
| scHiCSRS | 0.77(0.31) | 0.76(0.25) | 0.76(0.24) | 0.51(0.34) | 0.48(0.20) | 0.48(0.17) | 0.44(0.34) | 0.35(0.20) | 0.34(0.17) |
| MF | 1.02(0.92) | 1.02(0.92) | 1.02(0.92) | 0.89(1.03) | 0.90(1.03) | 0.89(1.03) | 0.81(0.96) | 0.82(0.96) | 0.82(0.97) |
| GK | 0.97(0.86) | 0.97(0.86) | 0.97(0.86) | 0.81(0.94) | 0.82(0.94) | 0.82(0.94) | 0.74(0.89) | 0.74(0.89) | 0.74(0.89) |
| RW | 1.01(1.23) | 1.01(1.23) | 1.01(1.23) | 0.90(1.32) | 0.90(1.33) | 0.90(1.33) | 0.96(1.30) | 0.96(1.30) | 0.96(1.30) |

(c) Type III

| | 7k | | | 4k | | | 2k | | |
|---|---|---|---|---|---|---|---|---|---|
| Methods | 10 | 50 | 100 | 10 | 50 | 100 | 10 | 50 | 100 |
| scHiCSRS | 0.42(0.28) | 0.39(0.18) | 0.39(0.15) | 0.34(0.27) | 0.26(0.16) | 0.25(0.13) | 0.32(0.28) | 0.21(0.15) | 0.19(0.12) |
| MF | 0.57(0.49) | 0.57(0.49) | 0.57(0.49) | 0.49(0.52) | 0.49(0.53) | 0.49(0.53) | 0.48(0.51) | 0.48(0.51) | 0.48(0.51) |
| GK | 0.54(0.47) | 0.54(0.47) | 0.54(0.47) | 0.45(0.48) | 0.45(0.49) | 0.45(0.49) | 0.44(0.48) | 0.44(0.48) | 0.44(0.48) |
| RW | 0.53(0.62) | 0.53(0.62) | 0.53(0.62) | 0.47(0.66) | 0.47(0.67) | 0.47(0.67) | 0.51(0.67) | 0.50(0.67) | 0.50(0.67) |

The numbers in the table are the average and those in the parentheses are the standard deviations over 100 replicates.

Table S7: Clustering results for three single-cell Hi-C data sets.

(a)GSE117876

| | GM | | PBMC | |
|---|---|---|---|---|
| Method | C1 | C2 | C1 | C2 |
| Observed | 13 | 1 | 7 | 11 |
| scHiCSRS | 13 | 1 | 5 | 13 |
| MF | 13 | 1 | 7 | 11 |
| GK | 13 | 1 | 7 | 11 |
| RW | 11 | 3 | 8 | 10 |

(b)GSE80006

| | K562A | | K562B | |
|---|---|---|---|---|
| Method | C1 | C2 | C1 | C2 |
| Observed | 1 | 1 | 0 | 8 |
| scHiCSRS | 2 | 0 | 0 | 8 |
| MF | 1 | 1 | 0 | 8 |
| GK | 1 | 1 | 0 | 8 |
| RW | 1 | 1 | 0 | 8 |

(c)scm3C-seq

| | L4 | | L5 | |
|---|---|---|---|---|
| Method | C1 | C2 | C1 | C2 |
| Observed | 76 | 55 | 105 | 75 |
| scHiCSRS | 131 | 0 | 6 | 174 |
| MF | 77 | 54 | 105 | 75 |
| GK | 77 | 54 | 104 | 16 |
| RW | 76 | 55 | 105 | 75 |

The results in the "Observed" are clustering results with observed data without imputation for data quality improvement.

Table S8: Computation time of the methods on three single cell Hi-C data sets.

| Method | GSE117874 | GSE80006 | scm3C-seq |
|---|---|---|---|
| scHiCSRS | 3.0m | 1h5m | 5.7h |
| MF | 0.8s | 19s | 5m |
| GK | 1.5s | 15s | 4m |
| RW | 0.1s | 4s | 2m |

#### References

K. Jin, L. Ou-Yang, X.-M. Zhao, H. Yan, and X.-F. Zhang. sctssr: gene expression recovery for single-cell rna sequencing using two-side sparse self-representation. *Bioinformatics*, 36 (10):3131–3138, 2020.

R. Vershynin. Introduction to the non-asymptotic analysis of random matrices. *arXiv preprint arXiv:1011.3027*, 2010.

Y. Zhao, Y.-J. Wu, E. Levina, and J. Zhu. Link prediction for partially observed networks. *Journal of Computational and Graphical Statistics*, 26(3):725–733, 2017.
